# An adaptive interaction between cell type and metabolism drives ploidy evolution in a wild yeast

**DOI:** 10.1101/2022.11.14.516370

**Authors:** Johnathan G. Crandall, Kaitlin J. Fisher, Trey K. Sato, Chris Todd Hittinger

## Abstract

Ploidy is an evolutionarily labile trait, and its variation across the tree of life has profound impacts on evolutionary trajectories and life histories. The immediate consequences and molecular causes of ploidy variation on organismal fitness are frequently less clear, although extreme mating type skews in some fungi hint at links between cell type and adaptive traits. Here we report an unusual recurrent ploidy reduction in replicate populations of the budding yeast *Saccharomyces eubayanus* experimentally evolved for improvement of a key metabolic trait, the ability to use maltose as a carbon source. We find that haploids have a substantial, but conditional, fitness advantage in the absence of other genetic variation. Using engineered genotypes that decouple the effects of ploidy and cell type, we show that increased fitness is primarily due to the distinct transcriptional program deployed by haploid-like cell types, with a significant but smaller contribution from absolute ploidy. The link between cell-type specification and the carbon metabolism adaptation can be traced to the noncanonical regulation of a maltose transporter by a haploid-specific gene. This study provides novel mechanistic insight into the molecular basis of an environment-cell type fitness interaction and illustrates how selection on traits unexpectedly linked to ploidy states or cell types can drive karyotypic evolution in fungi.

## INTRODUCTION

Ploidy is a fundamental aspect of the biology of all organisms, but it is subject to striking diversity across the tree of life—between related species, between individuals of the same species, and within individuals across cell types and life cycles [1]. The long-term impact of ploidy variation on eukaryotic evolution, particularly as a mechanism for generating raw material for natural selection, has long been recognized [2–6]. Recent work, primarily in the model eukaryote *Saccharomyces cerevisiae*, has further defined short-term evolutionary consequences of different ploidy states [7–13]. It remains less clear, however, what immediate effects on organismal fitness a ploidy transition can engender. In the diplontic species *S. cerevisiae*, ploidy variation is present both within the natural life cycle [14] and among isolates from diverse environments [15,16]. Despite this natural variation, diploidy seems to be generally favored [15]. Indeed, diploids frequently arise and sweep to fixation in laboratory evolution experiments founded with non-diploid strains [7,17–22].

In the limited cases where a direct fitness advantage of diploidy has been found in *S. cerevisiae* in the absence of confounding variation, the specific molecular basis or bases have remained elusive. A large-scale survey of *S. cerevisiae* and its sister species *Saccharomyces paradoxus* suggested that specific ploidy-by-environment interactions were necessary to explain observed differences in fitness proxies between haploids and diploids, which argues against generalizable predictions of fitness effects of ploidies across environments [23]. Similar experiments in *Candida albicans* found genetic background to influence fitness more than ploidy in several conditions that might be predicted to favor different ploidy states [24]. By contrast, more recent work capturing a wide swath of genetic diversity in *Saccharomyces eubayanus*, which diverged from *S. cerevisiae* ∼17 million years ago [25], failed to find meaningful differences in phenotypic traits between ploidies [26].

Adding complexity to the interpretation and prediction of fitness differences between ploidy states in yeast is the nuanced relationship between ploidy and cell type across species. In wild-type *Saccharomyces*, for example, ploidy indirectly controls cell type through the presence or absence of alleles at a single locus, the *<u>MAT</u>*ing type locus [14,27,28]. The haploid cell types express a common set of haploid-specific genes, as well as mating type-specific genes dependent on the allele present at the *MAT* locus, while diploids repress these gene sets but are competent to induce the expression of a small number of genes under specific conditions (e.g. meiosis). Although investigations into ploidy-specific fitness effects have primarily focused on the physiological differences between haploids and diploids that are independent of cell type, it remains plausible that underappreciated aspects of cell-type specification could influence traits that in turn impact organismal fitness.

Perhaps the most compelling evidence for widespread effects of selection on cell type across fungi can be found among pathogenic species. Highly skewed mating type ratios have been described among isolates of *Cryptococcus neoformans*, *Candida glabrata, Candida auris, Fusarium poae,* and *Fusarium verticillioides* [29–35]. Large mating type skews are also found in clinical isolates of *Aspergillus fumigatus* but not in isolates from other sources, and mating type has been shown to influence pathogenicity *in vitro* and virulence *in vivo* in this species [36,37]. Similar links between mating type and virulence traits have been suggested in *Cryptococcus neoformans*, *Candida auris*, *Mucor iregularis,* and *Fusarium graminearum* [38–46], suggesting that unexpected links between cell type and traits experiencing intense selection may be widespread among fungi.

Microbial traits and their underlying genotypes are of particular interest when they directly impact human health, are important for biotechnological processes, or serve as models of eukaryotic evolution. The latter two cases are exemplified in the emerging model yeast *Saccharomyces eubayanus,* the wild parent of hybrid lager-brewing yeast. Since its isolation as a pure species [47], *Saccharomyces eubayanus* has become a model for microbial population genomics and ecology [48–53], as well as a key target for applied biotechnological research [54–59]. A focal ecological and industrial trait in this wild species is the ability to consume and metabolize the α-glucoside maltose, which is the most abundant sugar in the wort used to brew beer [60,61]. This trait is nearly ubiquitous among isolates of *Saccharomyces eubayanus* and its sister species *Saccharomyces uvarum* [62], but it has been lost [63] or severely curtailed [64] in the Holarctic subpopulation of *S. eubayanus* [51]. Paradoxically, members of this group contain functional structural maltose metabolism genes, and their inability to use this sugar as a carbon source may be due to cis- or trans-regulatory evolution resulting in altered expression of structural metabolism genes.

In an effort to identify mechanisms by which maltose utilization might be refined or regained after secondary loss, we previously subjected a Holarctic *S. eubayanus* strain to adaptive laboratory evolution (ALE) under selection for improved growth on maltose [64]. Here we map the genetic basis of adaptation in the evolved clones. We find that, surprisingly, haploids emerged and rose to high frequency in replicate ALE populations founded with a diploid strain, which is a highly unusual ploidy transition for *Saccharomyces*. We find that haploidy confers a substantial fitness advantage in the ALE conditions, but that haploids experience a fitness tradeoff in rich conditions, consistent with previous observations in *S. cerevisiae*. We identify cell type as the primary driver of adaptive fitness, with a smaller but significant contribution from absolute ploidy. Finally, we demonstrate that a major fitness-modifying gene has elevated expression in haploids, and that this effect can be linked to unexpected regulation by a haploid-specific transcription factor that regulates invasive growth in *S. cerevisiae*. Our results mechanistically explain a ploidy-by-environment fitness effect and demonstrate how strong selection on traits linked to cell types can drive karyotypic evolution in fungi.

## MATERIALS AND METHODS

### Strains, plasmids, and cultivation conditions

Strains, oligonucleotides, and plasmids used in this work are listed in tables S1 and S2. Yeast strains were propagated on rich YPD medium (10g/L yeast extract, 20g/L peptone, 20g/L glucose, with 18g/L agar added for plates), Synthetic Complete medium with maltose or glucose (5g/L ammonium sulfate, 1.7g/L Yeast Nitrogen base, 2g/L drop out mix, 20g/L maltose or glucose, pH=5.8, with 18g/L agar added for plates), or Minimal Medium (5g/L ammonium sulfate, 1.7g/L Yeast Nitrogen base, 10g/L maltose or glucose, pH=5.8) at room temperature. Yeast strains and ALE populations were stored in 15% glycerol at −80° for long-term storage. For supplementation with drugs, 1g/L glutamic acid was substituted for ammonium sulfate in SC media. G418, Hygromycin B, and Nourseothricin (CloNAT) were added to media at final concentrations of 400mg/L, 300mg/L, and 50mg/L, respectively. Transformation of *S. eubayanus* was performed via a modified PEG-LiAc method [65] as previously described [64]. Repair templates for homologous recombination were generated by PCR using Phusion polymerase (NEB) and purified genomic DNA as template or Taq polymerase (NEB) and purified plasmid as template per the manufacturer’s instructions, followed by purification with QiaQuick or MinElute spin columns (Qiagen). For CRISPR-mediated transformations, pXIPHOS vectors [66] expressing Cas9 and a target-specific sgRNA were co-transformed into strains with double-stranded repair templates. Multi-fragment repair templates were assembled by overlap extension PCR with Phusion polymerase, or co-transformed as multiple linear fragments with 80bp overlapping homology for *in vivo* recombination. Following transformation, yeast cells were plated to YPD for recovery and replica-plated to medium containing the appropriate antibiotic for selection after 24-36 hours. Gene deletions and knock-ins were verified by colony PCR and Sanger sequencing.

Plasmids were propagated in *E. cloni* 10G cells (Lucigen) and purified using the ZR miniprep kit (Zymo Research). sgRNAs for CRISPR/Cas9-mediated engineering were designed using CRISpy-pop [67], obtained as single-stranded 60-mers from Integrated DNA Technologies, inserted into *Not*I-digested pXIPHOS vectors using HiFi assembly (NEB), and verified by Sanger sequencing.

### Growth assays

Strains were streaked to single colonies on solid YPD agar, and individual colonies were inoculated to 250μL YPD in flat-bottom 96-well plates for preculturing in a randomized layout. Precultures were incubated for three days at room temperature, serially diluted in Minimal Medium, and inoculated to Minimal Medium containing maltose or glucose at a final dilution factor of 10^-4^. Plates were incubated on a SPECTROstar Omega plate reader (BMG Labtech) equipped with a microplate stacker, and OD_600_ was measured every hour. Raw growth data was processed using GCAT [68] and further analyzed in R v4.0.4 (https://www.R-project.org) (Core R Team).

### *MAT* locus genotyping

We used a multiplex colony PCR with Taq polymerase (NEB) and oligos oHJC120, oHJC121, and oHJC122 to genotype the *MAT* locus of strains following tetrad dissection, mating type engineering, and for estimating the frequency of haploids in adaptive laboratory evolution populations after plating. The multiplex reaction gives rise to *MAT***a**- and *MAT***α**-specific amplicons of differing size, which were resolved on 2% agarose gels. All reaction conditions were per the manufacturer’s instructions and were carried out alongside controls (diploid *MAT***a**/*MAT***α**; haploid *MAT***a**; haploid *MAT***α**; no input DNA). We discarded any experiment where the controls did not produce the expected amplicons (or lack thereof). To estimate the frequency of haploids in populations, we screened a total of 55-56 single colonies across four independent platings of each population. We note that this approach cannot formally distinguish between cells of different ploidies with rare aberrant *MAT* locus composition (e.g. diploid *MAT***a**/*MAT***a** will generate the same amplicon pattern as haploid *MAT***a**; loss of *MAT* locus heterozygosity in diploid *S. cerevisiae* has been estimated to occur at a rate of 2×10^-5^ per cell per generation [20]). In addition, this *S. eubayanus* background is homothallic, meaning that any diploid colony recovered following plating might represent a haploid cell in the experimental population maintained in liquid medium. The rate of mating type switching and clone-mate selfing on solid medium is likely orders of magnitude higher than loss of *MAT* locus heterozygosity [14,70]; thus, our PCR-based estimates of haploid frequency may be conservative.

### Mating type testing

In addition to molecular validation of engineered strains, we tested the expressed mating type of strains with altered *MAT* locus composition using microbiological assays. To assess *MAT***α** expression, a saturated liquid culture of *S. cerevisiae bar1-Δ* was diluted 100-fold and spread-plated to YPD, and 10μL of overnight query strain culture was spotted on top. For *MAT***a** expression validation, saturated cultures of query strains were diluted 100-fold and spread-plated to YPD, and a disc of sterile filter paper saturated with 10μL of 200μM **α**-factor (Zymo Research) was gently embedded in the center. Every experiment included wild-type controls of known mating type (diploid *MAT***a**/*MAT***α**; haploid *MAT***a**; haploid *MAT***α**), and in each case, growth inhibition by **α**-factor or of *S. cerevisiae bar1-Δ* was scored relative to controls and compared to the parental strain, where applicable.

### DNA sequencing

To obtain high molecular weight genomic DNA from wild-type strain yHRVM108, two single colonies were each inoculated in 90mL YPD and grown to mid-log phase (OD_600_=0.5), harvested by centrifugation, washed with water, and resuspended in 5mL DTT buffer (1M sorbitol, 25mM EDTA, 50mM DTT). Cells were DTT-treated for 15 minutes at 30° with gentle agitation, pelleted, washed with 1M sorbitol, and resuspended in 1mL 1M sorbitol with 0.2mg 100T Zymolyase. Cells were spheroplasted for 30 minutes at 30° with gentle agitation, then pelleted. The pellet was gently resuspended in 450μL EB (Qiagen) without pipetting and treated with 50μL RNAse A (10μg/mL) for 2 hours at 37°. 55μL 10% SDS was added, and the mixture was incubated for a further hour at 37° with gentle agitation to lyse spheroplasts. DNA was extracted by the phenol/chloroform method and precipitated by addition of 1mL 100% ethanol and overnight incubation at −80°. Precipitated DNA was pelleted, washed twice with 70% ethanol, dried briefly, and gently resuspended in 100μL TE buffer at room temperature without pipetting for two hours. DNA was quantified using the Qubit dsDNA BR kit (Thermo Fisher Scientific), and purity was assessed by Nanodrop (Thermo Fisher Scientific).

DNA concentration was adjusted to 50ng/μL, and 7.5μg genomic DNA was subjected to SPRI size selection with Agencort AMPure XP beads in custom buffer following the recommended protocol from Oxford Nanopore Technologies. 1μg size-selected DNA was prepared for sequencing using the SQK-LSK109 ligation kit (Oxford Nanopore Technologies), and approximately 40fmol library was loaded on a single FLO-FLG001 flowcell. Basecalling was performed with Guppy v3.2.1. ONT sequencing yielded 885.1Mb of base-called reads passing quality filtering, for approximately 74-fold genomic coverage.

We prepared genomic DNA for Illumina sequencing from the wild-type strain and evolved isolates as described previously [25]. Strains were streaked to single colonies, and colonies were inoculated to 3mL YPD medium and grown to saturation before collection for DNA extraction. Purified DNA was quantified by Qubit dsDNA BR assay (Thermo Fisher Scientific), and purity and quality were assessed by Nanodrop (Thermo Fisher Scientific) and agarose gel electrophoresis. Library preparation and Illumina-sequencing were performed by the DOE Joint Genome Institute. Paired-end libraries were sequenced on a NovaSeq S4 with 150bp reads, yielding an average of 8.47 million reads per sample. For the wild-type strain yHRVM108, we also integrated publicly available reads (SRA: SRX1317977) from a previous study [49]. Raw reads were processed using Trimmomatic v0.3 [71] to remove adapter sequences and low-quality reads.

### Genome assembly, annotation, and analysis

Canu v1.9 [72] was used to generate a genome assembly with Nanopore sequencing reads from the wild-type strain, which was subsequently polished with Illumina reads using three rounds of Pilon v1.23 [73]. The genome assembly was annotated using the Yeast Genome Annotation Pipeline [74]. We mapped each predicted gene to its *S. cerevisiae* homolog using BLASTp v2.9 [75]. QUAST v5.0.2 [76] and BUSCO v3.1.0 [77] were used to assess genome completeness, and chromosomes in the assembly were assigned numbers corresponding to the *S. eubayanus* type strain reference genome [58,78] using MUMmer v3.2.3 [79] and BLASTn v2.9. BWA v0.7.12 [80] and samtools v1.9 [81] were used to map short reads from all sequenced strains to the assembly, and BEDtools v2.27 [82] was used to call sequencing depth. Coverage across the genome of each strain was analyzed in R and assessed by manually inspecting coverage plots of each chromosome. Final genome-wide Illumina-sequencing depths for each strain were 200.7-fold (wild-type), 106.1-fold (evolved clone 1), and 84.2-fold (evolved clone 2). We used FreeBayes v1.3.1 [83] to call variants in each strain, requiring a minimum coverage depth of 10 to report a position, and manually inspected putative variants in IGV [84]. To annotate predicted transcription factor binding sites in the promoter of *AGT1*, we used the 700bp upstream of the start codon as a query for YEASTRACT+ [85] and YeTFaSCo v1.02 [86] using the “expert curated—no dubious” motif set.

### RNA extraction, sequencing, and analysis

Strains were streaked to singles on solid YPD agar, colonies were precultured to saturation in synthetic complete medium with 2% glucose or maltose as the sole carbon source, and pre-cultures were back-diluted into the same medium at a low initial OD_600_ (≈0.05) before being harvested in mid-log phase (OD_600_ ≈0.6-0.8), a growth regimen designed to fully alleviate catabolite repression of alternative carbon metabolism genes. Each strain-carbon source combination was assayed in biological triplicate. Cells were harvested by centrifugation after the addition of 0.1 volumes of 5% acid phenol/95% ethanol, and pellets were flash-frozen. RNA was extracted using the hot acid phenol/ethanol precipitation method, but we added glass beads during vortexing to aid lysis efficiency. Genomic DNA was digested using Turbo DNAse (Promega), and RNA yield and quality were assessed by Qubit BR RNA assay (Thermo Fisher Scientific), agarose gel electrophoresis, and Qubit RNA IQ assay (Thermo Fisher Scientific).

mRNA enrichment, library preparation, and Illumina-sequencing were performed by the DOE Joint Genome Institute. Paired-end libraries were sequenced on a NovaSeq S4 with 150bp reads, yielding an average of 23.23 million reads per sample. Raw mRNA-seq reads were processed with BBduk (https://sourceforge.net/projects/bbmap/) to remove adapters and low-quality sequences, resulting in an average of 22.89 million surviving reads per library. Filtered reads were mapped to the wild-type strain assembly with HISAT2 v2.1 [87] with average mapping rates of 98.5% per sample. HTSeq-count v0.11.1 [88] was used to generate transcript counts at each gene in the annotation. Raw counts were passed to DESeq2 v1.30.1 [89,90] for further analysis. We removed from analysis a single library from an evolved isolate grown in maltose, as manual inspection of normalized gene expression values revealed that this sample had stochastically lost the ChrXV aneuploidy. This reduced our power to detect statistically significant differences in expression for that specific evolved isolate. All other samples from evolved isolates remained aneuploid in both conditions. We considered differentially expressed genes between conditions and genotypes with expression changes of greater than or equal to twofold in either direction and Benjamini-Hochberg adjusted *p*-values of less than or equal to 0.01 (false discovery rate of 1%). Full differential expression analysis results can be found in Table S4. To compare expression levels of single genes, we used size-normalized counts from DESeq2, which are more robust for this purpose than other normalization methods [91–93]. We defined subtelomeric genes as those falling within 20kb of the end of a contig, which represented entire chromosomes in our assembly (with the exception of the two-contig ChrXII, for which we considered genes within 20kb of the telomeric contig ends, not the ends containing *rDNA* repeats). This classification is comparable to or more conservative than those used previously [94,95]. GOrilla [96] was used to identify enriched gene ontology (GO) terms in gene sets of interest; we used *S. cerevisiae* GO annotations and specified all predicted genes in our annotation as the background set against which to test for enrichment. Statistics and data visualization were performed in R.

### Ploidy determination

Flow cytometry-based ploidy determination was performed as described previously [7], except that we sampled asynchronous cultures. Briefly, we fixed mid-log cultures of each query, treated fixed cells with RNAse A and Proteinase K, and stained DNA with Sytox Green (Thermo Fisher Scientific). Haploid and diploid *S. cerevisiae* strains were included in all experiments as controls. For clonal strains (*S. cerevisiae* controls, ancestral *S. eubayanus*, and evolved isolates), queries were streaked to single colonies, and independent colonies were picked for ploidy analysis. For population samples, entire populations cryopreserved at −80°C in 15% glycerol were gently thawed, and approximately 50μL of slurry was inoculated directly to 1mL SC-2% Maltose. The optical densities of these samples were monitored, and they were harvested and fixed in early log phase after a minimum of two doublings. We sampled 10,000 cells for each query on an Attune NxT flow cytometer (Thermo Fisher Scientific). Analysis was performed in FlowJo v10.

### Fitness assays

Except for experiments in rich medium shown in Fig. 2A, the conditions for fitness assays were designed to mimic the original ALE conditions [64]. Briefly, this regime consisted of culturing in 1mL SC medium with 2% maltose and 0.1% glucose (hereafter, “competition medium”) with semiweekly 1:10 dilutions into new competition medium. Query genotypes were directly competed against a common competitor in co-culture. The competitor was a haploid in the ancestral *S. eubayanus* strain background with the exception of a constitutively expressed GFP using a *TEF1* promoter and *ADH1* terminator from *Saccharomyces cerevisiae* and *ste12* deletion (*MAT***a** *hoΔ::P_ScTEF1_-yEGFP-T_ScADH1_-kanMX, ste12Δ::natMX*). We chose a *ste12* deletion to prevent any interaction with competitors expressing *MAT***α.** Strains were streaked to single colonies on YPD containing antibiotic as appropriate, precultured in competition medium for three days, mixed in approximately equal query-to-competitor ratios (except where we reduced the competitor ratio against less-fit query genotypes), sampled into cold 1X PBST for flow cytometry of timepoint 0, and inoculated into 1mL competition medium at an initial OD_600_ of approximately 0.1. At each transfer, competitions were sampled into cold 1X PBST for flow cytometry, and the optical density of each replicate was measured to calculate the number of generations. Competitions in rich medium were carried out in the same manner, albeit that preculturing and propagation were in sterile-filtered YPD in 2mL volume with daily dilutions of 1:100. For both competition regimes, we sampled 13,000 cells per replicate and timepoint on an Attune NxT flow cytometer (Thermo Fisher Scientific) to quantify the abundance of competitor (fluorescent) and query (non-fluorescent) cells, which always clearly formed distinct populations. Analysis was performed in FlowJo v10. Fitness was calculated as the selection coefficient, obtained by regressing the natural log ratio of query to competitor against the number of generations. To analyze the effects of ploidy, mating type, and cell type (diploid-like and haploid-like) on the panel of engineered strains shown in Fig. 3, we used multiple linear regression with measured fitness as the response and ploidy, mating type, and cell type as categorical predictors with two levels each (for mating type, we grouped by whether genotypes expressed any mating type-specific genes, or none). All statistical analyses and visualization were performed in R.

**Figure 1.**
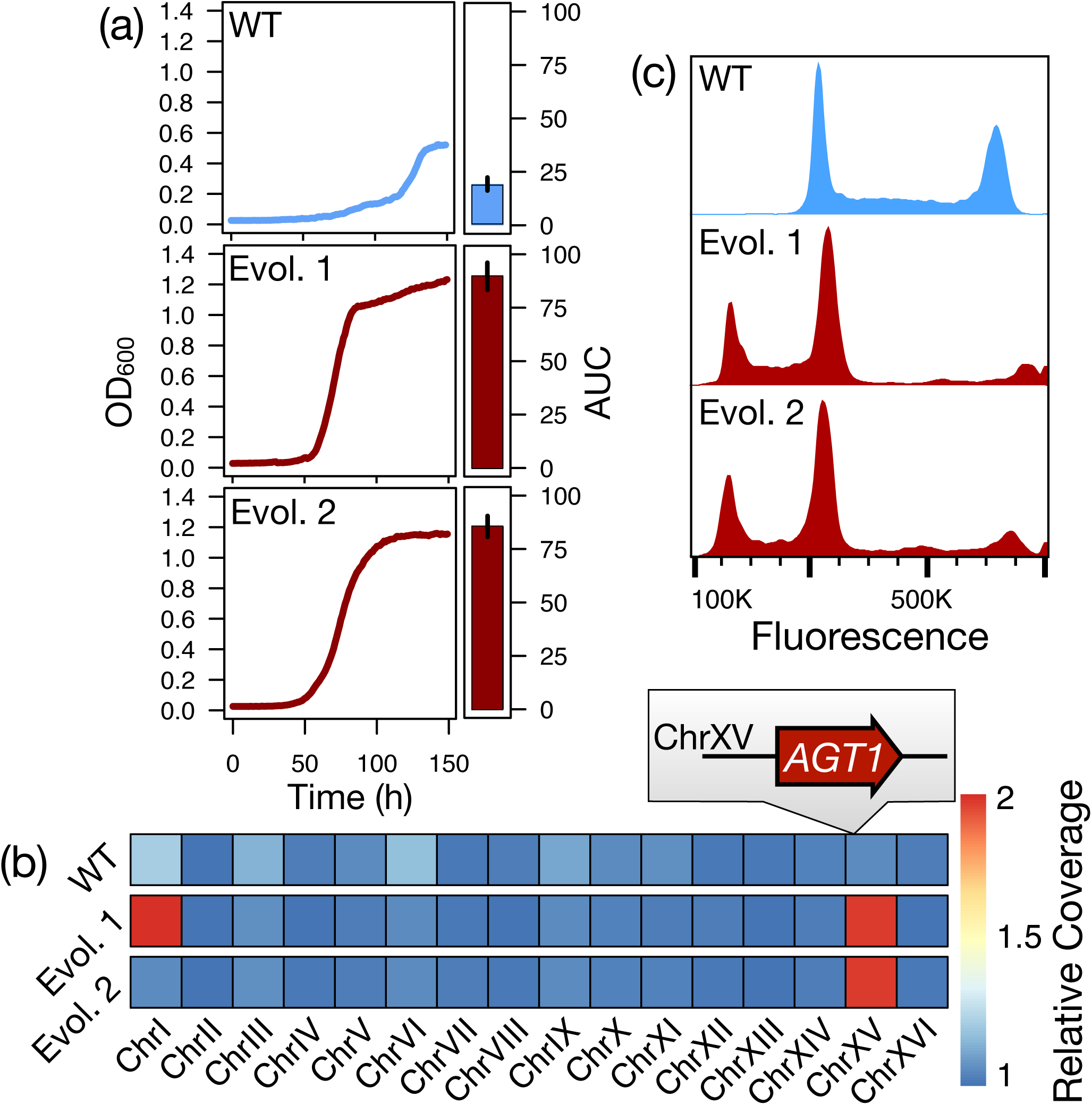
Phenotypic and karyotypic evolution of *Saccharomyces eubayanus* isolates. (a) Growth of the wild *S. eubayanus* strain (WT) and clonal isolates from two replicate experimental evolution populations (Evol. 1, Evol. 2) on maltose. A representative growth curve is shown for each; bar plots show mean and standard error of total growth (area under the curve, AUC) across six biological replicates for each genotype. (b) Relative copy number of each chromosome in the wild-type and evolved strains, inferred from sequencing depth. The parallel ChrXV gain includes a homolog of an *S. cerevisiae* gene encoding an α-glucoside transporter (*AGT1*/YGR289C). (c) Smoothed histograms of cellular DNA content in the WT and evolved strains as measured by flow cytometry. Primary peaks correspond to cells in G1 and G2, and fluorescence intensity is proportional to DNA content.

**Figure 2.**
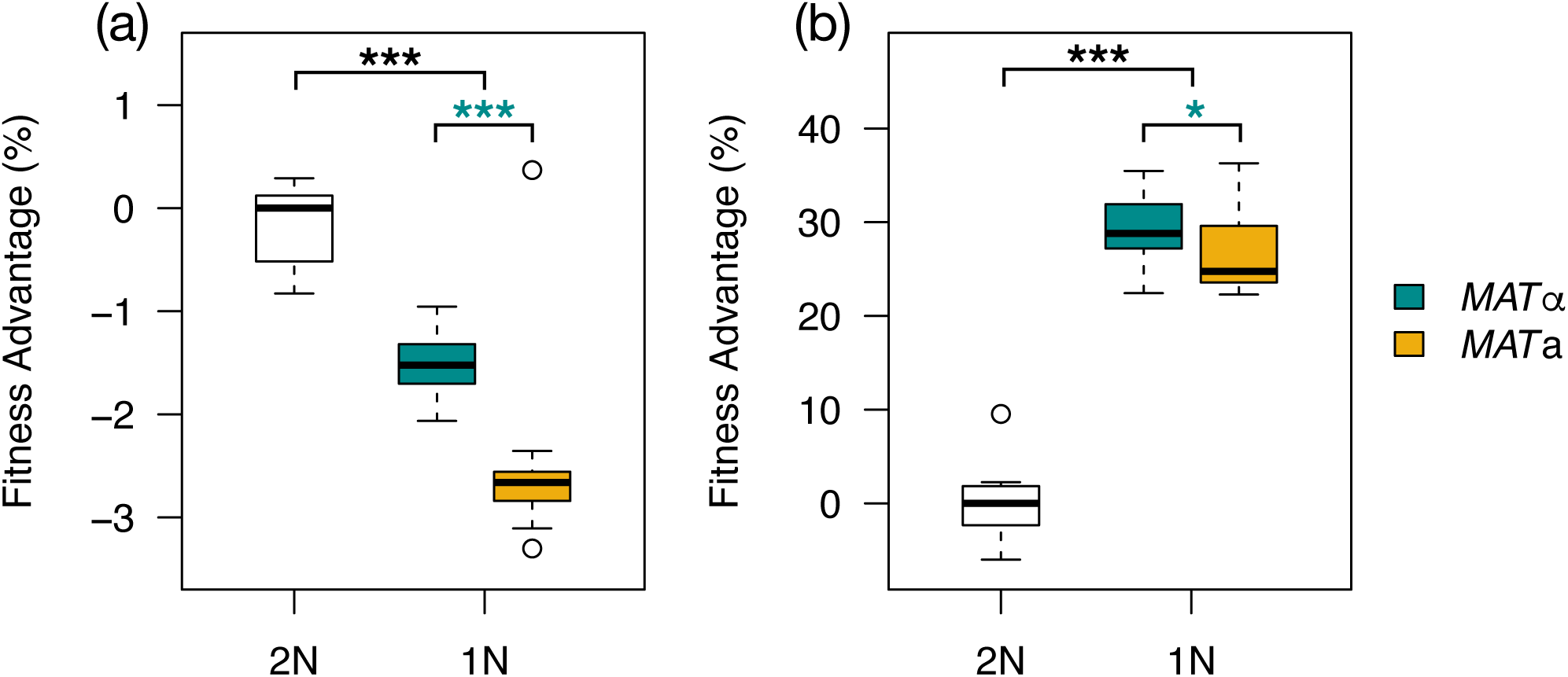
Haploids have a conditional fitness advantage. Boxplots show fitness measurements of isogenic diploids (n=12) and haploids from fully viable tetrads (n=48) in rich medium (a) and adaptive laboratory evolution (ALE) conditions (b). *** *p* < 10^-4^ (Mann-Whitney *U* tests) between diploids and each haploid group (black) or between haploid groups (teal). In ALE conditions, the significance level between haploid groups was 0.018 (*).

**Figure 3.**
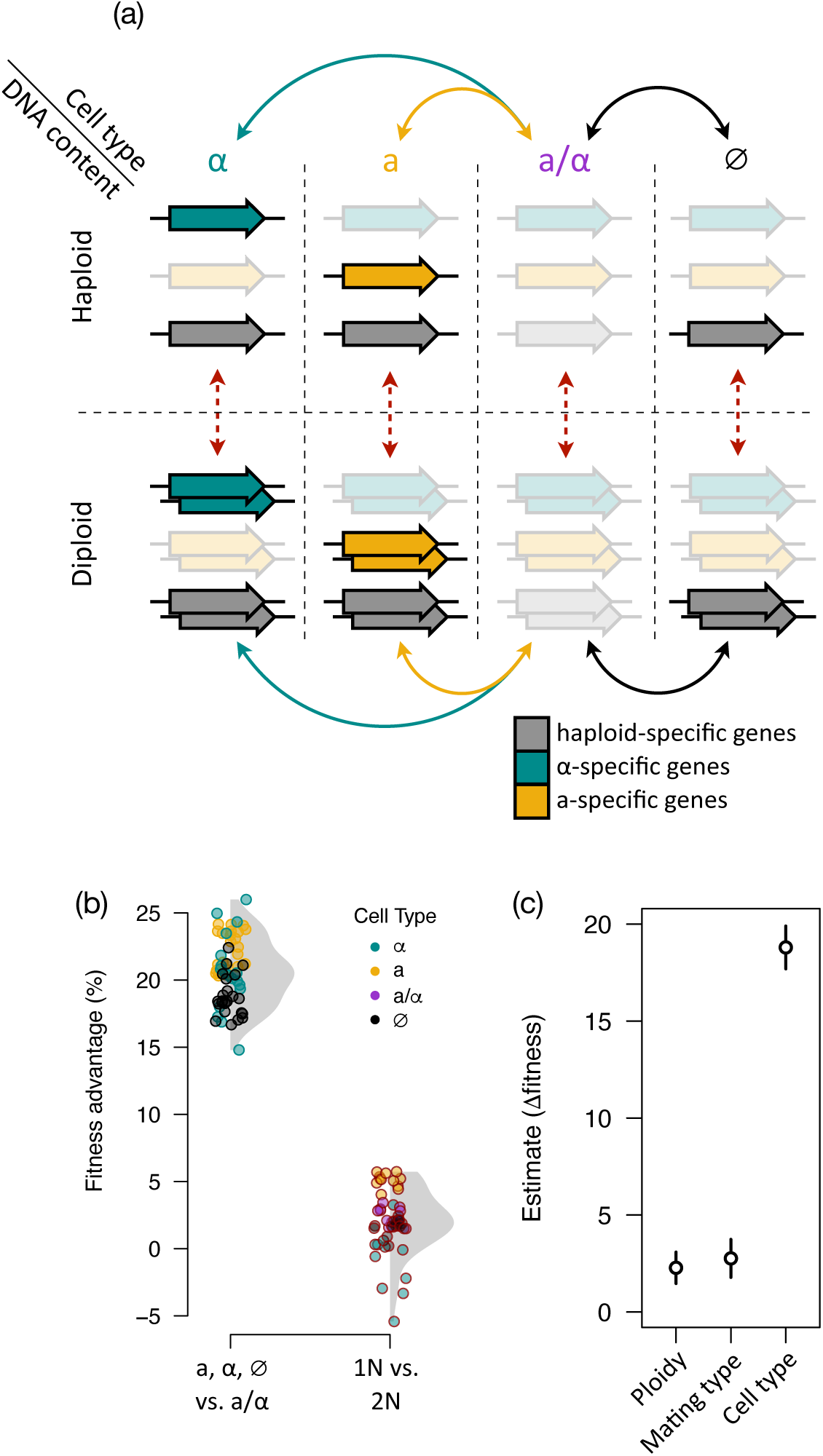
Cell-type specification is the primary contributor to adaptive fitness. (a) Schematic of engineered strains compared to determine ploidy and cell type effects on fitness. The three classes of cell type-specific genes are depicted as colored bars, with opacity indicating expression. Dotted red lines are comparisons that show the effect of absolute DNA content; solid lines are comparisons that show the effect of cell type. (b) Points show differences in fitness in ALE conditions between genotypes that differ in only cell type or ploidy. Grey shading shows the density distribution of each group. In each case, the wild-type state is taken as the baseline for comparison (diploid; **a**/**α** cell type). (c) Estimates and 95% confidence intervals for the effect of each variable on the change in organismal fitness.

### *P_AGT1_* reporter analysis

We generated single-copy genome integrations in haploids of yeast-optimized GFP (yeGFP) expressed from both the native *AGT1* promoter and a variant in which we abolished the Tec1p consensus site (TCS) by making point mutations to each of its six nucleotides. Sequence-verified GFP strains and controls were streaked to single colonies on YPD containing antibiotic as appropriate, picked to SC-2% maltose and grown to saturation, back-diluted in 2mL SC-2% maltose to an initial OD_600_ of 0.01, and grown to mid-log phase (OD_600_ ≈0.5-0.7). Cells were collected by centrifugation, washed twice with cold PBST, and resuspended in PBST for flow cytometry. We sampled 40,000 cells per replicate on an Attune NxT (Thermo Fisher Scientific). Analysis was performed in FlowJo v10. Fluorescence values were exported for statistical analysis and visualization in R.

## RESULTS & DISCUSSION

### Evolved *S. eubayanus* isolates harbor mutations incongruous with ancestral ploidy

We previously experimentally evolved a wild strain of *Saccharomyces eubayanus* from the Holarctic subpopulation under selection for improved growth on the industrially relevant α-glucoside maltose [64]. We picked clones from two replicate populations of the ALE experiment that displayed significantly increased growth (*p*=0.002, Mann-Whitney *U* tests) on maltose compared to the ancestral strain (Fig. 1A). To map the genetic basis of improved growth on maltose, we sequenced the genomes of each clone to a final average depth of 95-fold. We mapped these reads to a re-sequenced and annotated assembly of the ancestral strain and identified a total of four single nucleotide polymorphisms (SNPs) and three large-scale copy number variants (CNVs) in the form of aneuploidies across the evolved isolates (Fig. 1B, Table S3). We did not identify single-nucleotide variants in or near any genes with clear relationships to α-glucoside metabolism, although one SNP introduced a premature stop codon in *IRA1*, a common target of adaptive mutations in batch-style experimental evolution [7,22,97–99]. One aneuploidy (ChrXV gain) was shared between evolved isolates and encompassed a homolog of the *S. cerevisiae* generalist α-glucoside transporter *AGT1*/YGR289C, suggesting a potential mechanism for adaptation (Fig. 1B).

We noted two unexpected features of evolved mutations: all SNPs in the evolved isolates were represented by a single, non-reference allele (Fig. S1), and all aneuploidies identified were present at twofold relative copy number, while sequencing depth across the genome of the ancestral strain indicated euploidy. Although mitotic recombination can generate losses of heterozygosity at new or standing variation during adaptive evolution [100–105], our results differed significantly from two recent large-scale experimental evolution studies in *S. cerevisiae*, which found approximately 5-10% of mutations to be homozygous in diploid or autodiploid clones after 4,000 generations [7,9]. In comparison, our observed allele frequencies at mutated sites are highly improbable under the null expectation of diploidy (binomial tests: *p*=5.3×10^-6^, *p*=1×10^-4^, respectively). Similarly, while aneuploidies are common in both wild *Saccharomyces* isolates and as outcomes of experimental evolution or mutation accumulation, tetrasomies—resulting in twofold relative copy number—are relatively rare in diploids [8,15]. Thus, we reasoned that the observed patterns in copy number and allele frequency might best be explained by an unexpected and atypical ploidy reduction to haploidy during ALE.

### Haploids emerged and rose to high frequency in diploid-founded populations

We directly determined the ploidy states of the evolved clones and the ancestral strain using flow cytometry (Fig. 1C) and confirmed that the strain that was used to found the experimental populations was diploid. Consistent with the results of genome sequencing, we found that clones from both ALE replicates had become haploid. To test whether the clonal isolates we analyzed were simply from a rare and non-representative subpopulation, we assayed the ploidy states present at the population level in both replicates of the ALE experiment (Fig. S2B). Haploids were clearly detectable in each replicate by generation 100 and 250, respectively. As an orthogonal approach, we plated cells from the terminal timepoint of each population of the ALE experiment and used a PCR assay to genotype the *MAT* locus of single colonies. By this method, haploids constituted 74-100% of the cells we genotyped in the two ALE populations (Fig. S2C). Thus, although haploids did not sweep to fixation in both experimental populations, they repeatedly emerged and rose to high frequency over the duration of the ALE experiment.

### Haploids exhibit a direct condition-dependent fitness advantage

The abundance of haploids in our experimental populations could be explained by two alternative models: haploids might have a direct fitness advantage, or they might benefit indirectly from increased adaptability in our ALE environment. Two well-documented lines of evidence from previous studies seemed to strongly favor the latter hypothesis. First, *S. cerevisiae* haploids have repeatedly been shown to adapt more rapidly than diploids during experimental evolution, in part due to dominance effects at adaptive targets and ploidy-specific mutation rates and spectra [7–12,106,107]. The common mutation shared by our adapted clones, ChrXV aneuploidy, might be predicted to have a larger effect size in haploids than diploids due to the difference in relative copy number conferred by the gain of a single chromosome copy between ploidies and the general concordance between increased copy number and gene expression in yeast [108–113]. Second, *S. cerevisiae* displays a strong trend of converging on a diploid state during experimental evolution initiated with non-diploid strains [18]. Although theory predicts that haploids may be better able to meet their metabolic needs in nutrient-limiting conditions due to increased cell surface area-to-volume ratios, experimental evidence in yeast has failed to find widespread support for such generalizable trends [23,114–117], and our experimental evolution conditions could not strictly be considered to be limited in key nutrients. Given the relative simplicity of testing for differences in fitness between ploidies, we first sought to support or refute the model of direct haploid advantage.

We used a sensitive competition assay to measure the competitive fitness of isogenic diploids and haploids in the wild strain background following *HO* deletion, sporulation, and tetrad dissection. Consistent with observations in *S. cerevisiae* of direct or cryptic diploid advantage [7,15,17,18,20,22,118], haploids in our strain background exhibited median fitness defects of 1.5% (*p*=1.5×10^-5^) to 2.7% (*p*=9.9×10^-5^) relative to the isogenic diploid in rich medium (Fig. 2A). By contrast, in the ALE conditions, haploids displayed median fitness advantages of 24.8% (*p*=1.6×10^-9^) to 28.8% (*p*=3.6×10^-9^) per generation over diploids (Fig. 2B). Interestingly, we observed a significant fitness difference between haploids of opposite mating types in both environments tested (rich medium *p*=1.1×10^-9^, evolution conditions *p*=0.018), suggesting a common underlying mechanism linked to mating type, rather than a specific mating type-by-environment interaction. Expression of the mating-type genes is known to be costly [119], making components of this pathway common targets of adaptive loss-of-function mutations in haploids [7,98]. The observed fitness defect of *MAT***a** haploids in our experiments may reflect an expression burden imposed by the greater number of *MAT***a**-specific genes; a metabolic burden imposed by synthesizing the more complex, post-translationally modified **a**-factor pheromone; or both. While previous large-scale studies in *S. cerevisiae, S. paradoxus,* and *S. eubayanus* have not reported general fitness differences between mating types of otherwise isogenic haploids [23,26], the subtle, but significant, differences we observed here may have been below previous limits of detection. Irrespective of mating type, we find that haploids have a large and unexpected advantage over diploids under the ALE conditions.

### Haploid fitness advantage is primarily due to cell-type specification

In *Saccharomyces*, ploidy is intrinsically linked with cell- and mating-type specification, which are determined by the allelic composition of the *MAT* locus [28]. Some differences in cell physiology and gene expression patterns between ploidies are attributable solely to total cellular DNA content, while loss of heterozygosity at the *MAT* locus establishes one of two partially overlapping, cell type-specific gene expression programs [27,120,121]. The relationship between DNA content and cell-type specification can serve to confound inferences of the underlying basis of fitness differences between ploidies, although in limited cases, contributions of either absolute ploidy or *MAT* locus composition have been documented [7,23]. Here we refer to “cell-type specification” as the distinction between genotypes with a full complement of cell-type master regulators (e.g. wild-type diploids containing *MATa1*, *MATα1*, *MATα2*) and those without. Cell types established by the absence of one or more cell-type regulators (e.g. wild-type haploids) effect the de-repression of a handful of genes, commonly referred to as “haploid-specific,” but whose expression is technically independent of ploidy and mating type.

To dissect the contributions of DNA content and cell type to organismal fitness in our system, we engineered a series of eight otherwise isogenic genotypes with unique combinations of ploidy, mating type, and cell type-specific gene expression. Beginning with a diploid *ho* deletion in the wild-type background and haploids of each mating type, we manipulated *MAT* gene composition to generate diploids specifying a haploid-like cell type and a single mating type (by deleting one copy of the *MAT* locus at random) or no mating type (by deleting *MATα1* in a *MATa-Δ* background), as well as in haploids specifying no mating type (by deleting *MATα1*) or a diploid-like cell type (by integrating a copy of the other *MAT* allele in a haploid of the opposite mating type). We measured the fitness of these strains in the ALE condition and estimated the separable effects of ploidy, mating-type specification, and cell type-specific gene expression patterns on fitness (Fig. 3). These three factors explained the majority of the variance in measured fitness across genotypes (multiple R^2^=0.96, df=86, *p* < 2.2×10^-16^), with each having a significant effect (*p* ≤ 2.56×10^-7^). Remarkably, cell-type specification had an impact on organismal fitness that was almost an order of magnitude greater than either ploidy or mating type (Fig. 3B, fitness advantage 18.8%, 95% CI: 17.7, 19.9), and explained far more of the variance (proportion sum squares: cell type, 0.93; ploidy, 0.016; mating type, 0.014). Absolute ploidy nonetheless impacted fitness across cell types, with haploids experiencing a 2.3% advantage relative to diploids in the ALE condition (95% CI: 1.5, 3.1). Paradoxically, expression of mating type-specific genes appeared to modestly increase fitness between haploid-like cell types in the ALE condition, in contrast to the documented cost of their expression in other conditions [119]. While one possible interpretation is that both sets of mating type-specific genes confer bone fide fitness advantages to cells growing in maltose medium, a more likely explanation for this apparent discrepancy is that haploid-like, *MAT*-null cells experience modest off-target fitness defects as a result of their extensive genetic engineering in the stringent ALE condition. As such, our analyses may slightly underestimate the fitness benefit attributable to haploid-like cell type in the ALE condition. We conclude that the cell type specified by the *MAT* locus, rather than absolute ploidy per se, has the largest effect on fitness in the ALE condition.

### A fitness-modifying maltose metabolism gene has cell type-specific increases in expression

The conditional fitness advantage of haploids and increased fitness of haploid-like cell types suggested an unexpected regulatory link between maltose metabolism and haploid-specific genes (i.e. those genes de-repressed in the absence of a heterozygous *MAT* locus). To identify potential targets of this interaction, we analyzed mRNA-seq data collected from the wild-type diploid and evolved haploids grown in conditions mimicking the evolution experiment (SC-maltose), as well as a baseline for comparisons (SC-glucose). Although the haploid strains had discrete polymorphisms, they shared a common cell type and aneuploidy of chromosome XV (Fig. 1B); thus, we reasoned that common differences in expression between these isolates relative to the wild-type strain should be attributable to one (or both) of these shared genotypes. Transcriptomes of the evolved haploids were highly similar, as expected (ρ=0.85, Fig. S4B), with genotype accounting for 18% of the total variance in gene expression across all samples in principal component space (Fig. S4A). Differentially expressed genes (DEGs) between the wild-type strain and evolved haploids were enriched for cell- and mating-type specific transcripts and genes on aneuploid chromosomes; however, there was no clear functional enrichment among DEGs to explain the maltose-specific haploid fitness advantage. The *AGT1* transporter on ChrXV was the single maltose metabolism-associated gene upregulated in maltose in both evolved haploids when compared to the wild-type strain, which was expected given its twofold relative copy number in these isolates (Fig. 1B). Upon closer examination, however, *AGT1* expression was higher than the twofold increase expected commensurate with its relative copy number [122,123]. Indeed, *AGT1* expression in haploids exceeded null expectations based on two distinct models (Fig. 4A): 1) we calculated the fold-change for *AGT1* in the ancestral strain in maltose compared to glucose and applied this multiplier to the glucose expression level in the evolved haploids; 2) we applied a twofold multiplier to the gene expression levels in the wild-type strain in both glucose and maltose, which accounted for copy number variation in the evolved haploids. While *AGT1* expression in glucose in the evolved haploids was in line with the naïve aneuploid expectation (*p*=0.81, one-sided Mann-Whitney *U* test), its expression in maltose in the evolved haploids was an average of 76% higher than could be modeled by accounting for copy number and native regulation (*p*=0.0005, one-sided Mann-Whitney *U* test).

**Figure 4.**
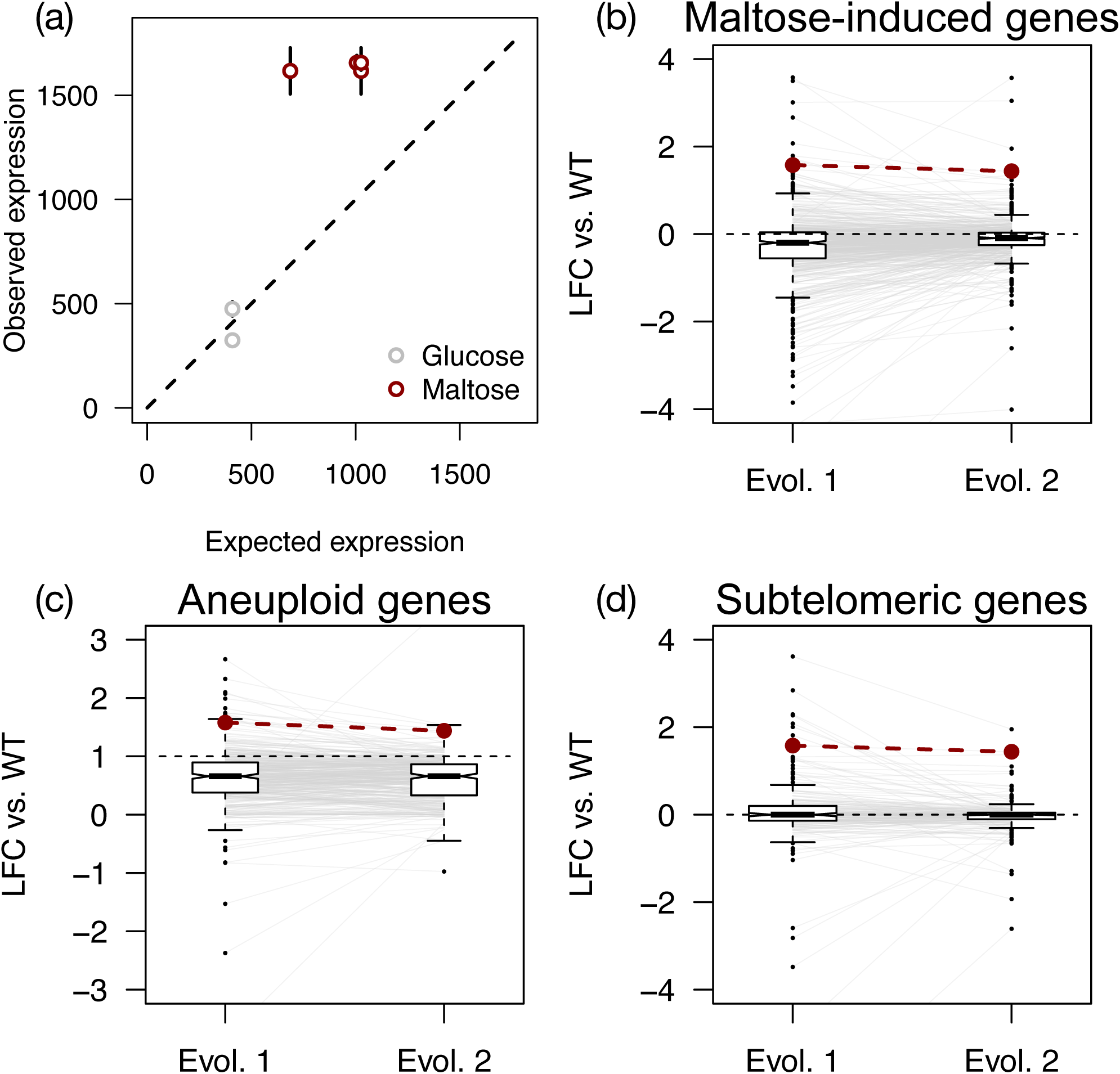
Increased expression of an α-glucoside transporter gene in haploids. (a) Points and bars show mean and standard error of *AGT1* expression in evolved haploids, plotted against the null expectation of expression based on copy number variation and induction in the wild-type strain. Expression in maltose is greater than the null expectations in the evolved haploids (*p*=0.0005, one-sided Mann-Whitney *U* test). (b-d) Boxplots show log_2_-transformed fold changes (LFC) of gene expression in maltose in evolved haploids compared to the wild-type (WT) strain. Whiskers extend to 1.5X the interquartile range. Lines connect the y-axis coordinates of the same gene in each evolved isolate; axes are scaled such that an occasional outlier is truncated from the plot space for a single strain. *AGT1* expression is plotted as red dots and lines, and black dashed lines indicate the null expectation for expression values. (b) Genes induced in maltose in the wild-type strain (n=544). (c) Genes on aneuploid ChrXV (n=370). (d) Subtelomeric genes (n=200). For all classes, expression in either evolved haploid is not significantly greater than the null expectation (one-sided t-tests, min. *p*=0.42).

We next asked whether increased *AGT1* expression could be explained by subtle changes in global gene expression levels between the wild-type diploid and evolved haploids. We compared expression levels of two relevant classes of genes under which *AGT1* falls and which we reasoned might be subject to modest differential expression: maltose-induced genes and subtelomeric genes (Fig. 4B, Fig. 4D). We also examined expression of genes on the aneuploid ChrXV to test whether these broadly exceeded the expectation of a twofold expression increase commensurate with copy number (Fig. 4C). In each case, expression in the evolved haploids was not significantly greater than the null expectation (one-sided t-tests, *p* > 0.4), and *AGT1* was in the upper tail of gene expression values for each class. Most notably, *AGT1* expression in maltose ranked higher than 95.9% and 98.6% of other aneuploid genes in each evolved haploid, respectively.

Our expression analyses are consistent with the observation that DNA copy number and mRNA abundance are broadly correlated, if not always directly proportional, in yeasts [108–111,113,122]. Median expression levels of 1.37—2.02-fold have been reported for systematically aneuploidized chromosomes present at a twofold relative copy number in otherwise isogenic strains of *Saccharomyces cerevisiae* [112]; in the same study, elevated expression of aneuploid genes also became attenuated during experimental evolution, analogous to the history of the adapted clones we sequenced here. Compared to the wild-type euploid, we observed median fold-changes for ChrXV genes of 1.58 in maltose (for both evolved haploids) and 1.72 and 1.70 in glucose, with potential confounding influences of cell type and attenuation following evolution. It remains formally possible that a fraction of the cells in culture stochastically lost the ChrXV aneuploidy during growth, although its maintenance in presumably non-selective conditions (SC-glucose for mRNA-seq; YPD for WGS) in both strains argues against this possibility. Indeed, the effect of cell type on *AGT1* expression becomes even more evident in light of the median expression levels of aneuploid genes in haploids: *AGT1* is upregulated an average of 4-fold in maltose across the haploid strains, while median fold-changes for all ChrXV genes between maltose and glucose are 0.969 and 0.970 for each haploid, respectively.

We next considered whether increased expression of *AGT1* alone contributes to overall fitness. Agt1p is a homolog of the well-characterized *S. cerevisiae* α-glucoside transporter, but in contrast to canonical *MAL* gene clusters that contain structural and regulatory maltose metabolism genes, *S. eubayanus AGT1* is isolated in the subtelomeric region of ChrXV. In our genome assembly, no predicted genes intersperse the *AGT1* start codon and the beginning of telomeric repeats some 6,770bp upstream. We did not identify any homologs of genes encoding *MAL* regulators, transporters, α-glucosidases, or isomaltases on ChrXV, nor was there any significant enrichment of functional categories for genes on this chromosome. As transport is generally regarded as the rate-limiting step for carbon metabolism in yeasts [124–127], even modest increases in transporter expression level could influence phenotype sufficiently for selection to act; indeed, regulatory variants with smaller effects on gene expression show signatures of natural selection in other eukaryotic systems [128–131]. To test whether increasing the expression of *AGT1* affects fitness, we inserted an additional copy of *AGT1* under its native promoter and terminator into the genomes of diploids and haploids at a separate site, and we measured the fitness of the resulting strains in the ALE conditions. Increased *AGT1* copy number conferred a substantial and significant fitness benefit across backgrounds (Fig. S3). Haploids received a more modest increase in fitness than diploids, likely attributable to diminishing returns epistasis, but they were significantly more fit overall. There was no interaction between the haploid mating type and the fitness effect of increased *AGT1* expression (*p*=0.8, Mann-Whitney *U* test). The increase of *AGT1* expression that we observed in haploids is therefore likely to affect phenotype in an adaptive manner.

### The *AGT1* promoter integrates cell-type and sugar-responsive regulatory networks

We investigated potential regulators of *AGT1* by scanning its promoter for putative transcription factor-binding sites using high-confidence *S. cerevisiae* motifs. This analysis identified binding motifs for two canonical regulators of maltose metabolism genes, Mal63p and Mig1p (Fig. 5A), in an organization consistent with the characterized regulatory module that controls the expression of maltose metabolism genes in *S. cerevisiae* [132–134]. Although a causal relationship has not been directly established, the presence of Mal63p consensus sequences upstream of maltose metabolism genes is well correlated with their induction by maltose in the type strain of *S. eubayanus* [58]. Predicted binding sites in the *AGT1* promoter were also unexpectedly enriched for transcription factors regulating filamentous (GO:0010570; FDR: 8.12×10^-9^) and pseudohyphal growth (GO:2000220, FDR: 2.7×10^-5^), as well as response to starvation (GO:0042594; FDR: 6.68×10^-3^). The former two categories are particularly noteworthy because these pathways have regulatory cross-talk with cell type-specific gene expression programs and can link alternative regulatory patterns to extracellular nutrient levels [135,136]; in particular, filamentous growth can be induced in response to depletion of fermentable carbon sources [137,138]. Indeed, a haploid-specific gene, *TEC1*, is required for filamentous growth [27,139,140]. Although it is best characterized for its cooperative interaction with Ste12p, Tec1p can activate target genes as a monomer in a dosage-dependent fashion [141–143], and it has been experimentally mapped to the Tec1p consensus sequence (TCS) in vivo across the genus *Saccharomyces* [144]. We identified a predicted Tec1p-binding site in the promoter of *AGT1* with 100% identity to the TCS (Fig. 5A) and hypothesized that Tec1p could mediate the cell type-specific increase in *AGT1* expression we observed in haploids.

**Figure 5.**
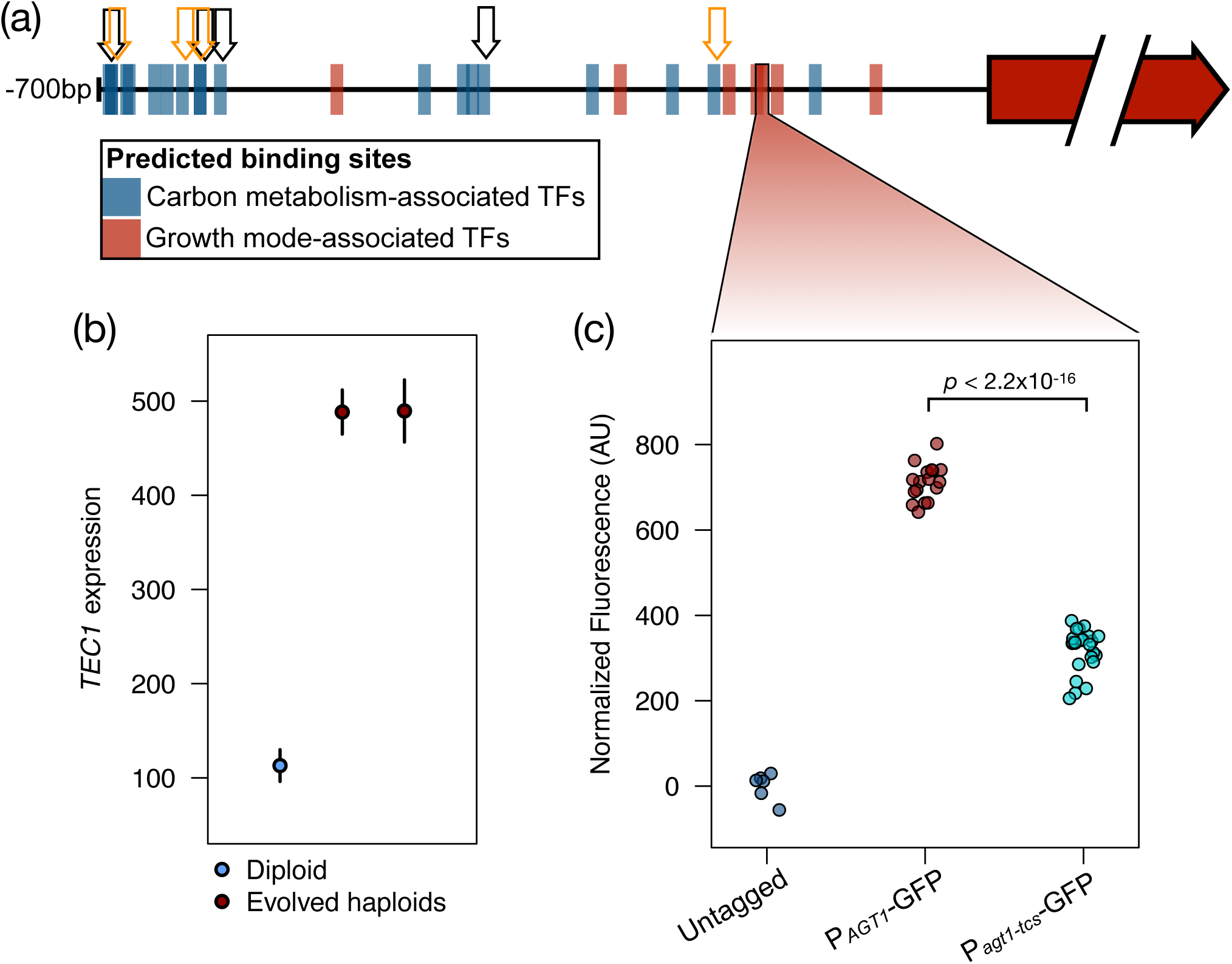
*AGT1* is regulated by cell type. (a) Predicted transcription factor-(TF-) binding sites in the promoter of *AGT1* for TFs related to carbon metabolism or growth mode. Boxes represent putative TF-binding sites in the 700bp upstream of the *AGT1* start codon and are colored by the functional characterization of the *S. cerevisiae* homolog. Arrows mark the predicted binding sites of two transcription factors that control *MAL* gene expression in *S. cerevisiae*, Mal63p (orange) and Mig1p (black). The position of a Tec1p consensus sequence (TCS) motif is marked with a black box. (b) *TEC1* expression is cell type-dependent in *S. eubayanus.* Points and bars show mean and standard error of *TEC1* expression (normalized counts) in the wild-type diploid and evolved haploids, averaged across conditions. (c) Point mutations to the predicted Tec1p-binding site in the *AGT1* promoter reduce reporter expression. Each point shows the mean population fluorescence for a replicate experiment with a control untagged strain (grey), as well as strains expressing GFP from the wild-type *AGT1* promoter (red) or a promoter with a mutated Tec1p binding site (teal). All engineered strains are significantly different from the untagged control (*p* ≤ 4.3×10^-6^, two-sided t-tests), and groups of promoter genotypes differ significantly (two-sided t-test).

To test this hypothesis, we generated single-copy genomic integrations in haploids of a GFP construct under the control of the wild-type *AGT1* promoter (*P_AGT1_*), as well as a promoter variant with point mutations in the predicted Tec1p-binding site (*P_agt1-tcs_*). We then measured single-cell fluorescence of the resulting strains grown in maltose by flow cytometry. Mutation of the Tec1p-binding site significantly decreased fluorescence from the reporter construct compared to the wild-type promoter (*p* < 2.2×10^-16^, two-sided t-test), but it did not abolish expression completely (Fig. 5C). These results are consistent with the expression data and collectively suggest that *AGT1* receives regulatory input from both cell-type and sugar-responsive networks, with separable activation by Tec1p and induction in the presence of maltose. In synthesis, the evidence for a direct fitness advantage by haploid-like cell types, increased expression of fitness-modifying *AGT1* in haploids, and the dependence of *AGT1* expression on the motif for a haploid-specific transcription factor paints a clear picture of a mechanistic relationship between ploidy evolution and adaptation in our system.

## CONCLUSIONS

Resolving the genotype-to-phenotype map remains a central goal in genetics and evolutionary biology, but it has frequently proven challenging, even in microbes. While gene content is generally correlated with metabolic traits across budding yeasts [25], regulatory nuances in organisms that are not traditional models can confound superficial inferences of phenotypes from genotypes [63,66,145]. In the taxonomic type strain of *S. eubayanus*, structural maltose metabolism genes in canonical *MAL* clusters are exquisitely repressed or induced hundreds-fold in response to carbon source [58], which is similar to their *S. cerevisiae* homologs [146]. By contrast, in the strain from the Holarctic subpopulation studied here, what appears to be the focal maltose transporter is partially decoupled from such stringent catabolite regulation: *AGT1* is only induced ∼2.3-fold in the wild-type strain in maltose (Table S4). We can envision two potential explanations for the apparently unusual regulation of this gene.

First, *AGT1* is likely to encode a transporter with broad substrate affinity like its *S. cerevisiae* homolog [64,147–150], whereas other phylogenetically distinct maltose transporters tend to have higher specificity [94,124]. It is possible that selection favored placing control of this generalist transporter under a broader transcriptional response to starvation or glucose depletion as part of a scavenging strategy, which the transition to filamentous growth is thought to represent [137]. Decoupling alternative carbon metabolism genes from their stringent canonical regulation has been shown to be adaptive among isolates of *S. cerevisiae* subject to specific ecologies [151] and ALE in fluctuating environments [152].

Second, the specific organization of *AGT1* in the *MAL1* locus of model *S. cerevisiae* strains—and its resulting exquisite regulation by glucose and maltose—could itself represent a derived state that is not reflective of wild yeasts. In strains of *Saccharomyces paradoxus, Saccharomyces mikatae,* and *S. eubayanus*, *AGT1* homologs are scattered in subtelomeric regions and not in canonical *MAL* loci, while homologs encoding high-affinity maltose transporters tend to occur in gene clusters with the typical organization [63,133]. Indeed, there is clade-specific variation within *S. cerevisiae* as to whether the *MAL1* locus is occupied by the generalist *AGT1* or a gene encoding a high-specificity maltose transporter [153], suggesting that domestication may have shaped the genetic architecture of α-glucoside metabolism in this model eukaryote [154]. Supporting this notion, *AGT1* homologs can be readily detected in publicly available *Saccharomyces* genomes, while growth on maltotriose—a sugar transported by *AGT1* but not most other maltose transporters—is extremely rare [62]. A notable exception is *Saccharomyces jurei,* the first wild *Saccharomyces* reported to grow on maltotriose, which contains a clear homolog of *AGT1* that requires extensive starvation or depletion of fermentable carbon sources for its induction [155]. Whether the organization and regulation of *AGT1* in Holarctic *S. eubayanus* represent a derived or ancestral state, it created a paradigm wherein a transition between ploidy states—and thereby cell types—was the adaptive step conferring the greatest increase in fitness among evolved genotypes we tested.

Remarkably, there is a strong parallel to the rewiring of carbon metabolism to cell type control in certain domesticated strains of *Saccharomyces cerevisiae*. Diastatic strains (sometimes called *Saccharomyces cerevisiae* var. *diasticus*) are characterized by their hyperattenuation, which is attributable to the presence of a novel extracellular glucoamylase, encoded by *STA1* [156]*. STA1* is a chimeric gene, created by the fusion of the sporulation-specific intracellular glucoamylase gene *SGA1* with the promoter and portions of the coding sequence of *FLO11*, which encodes a flocculin involved in filamentous growth that is subject to cell type-specific regulation [157–162]. Due to this gene fusion, *STA1* is expressed in a cell type-specific manner, and its regulation integrates catabolite repression by glucose and direct activation by Tec1p [163–165]. The cell type-dependence mediated by Tec1p in this case may have caused selection for haploidy among diastatic strains of the Beer 2 clade of *Saccharomyces cerevisiae* [15,166], which lack the clade-specific *AGT1* allele at the *MAL1* locus [153] and therefore must hydrolyze higher-order maltodextrins extracellularly.

*STA1* in diastatic brewing strains, *AGT1* in our ALE strains, and genes related to pathogenesis across fungi have undoubtedly experienced intense bouts of selection, and it seems that ploidy and cell type changes may be a common means of adapting, at least in microbial eukaryotes that have this flexibility. Here we have shown that the integration of regulatory inputs from metabolism and cell-type networks by the *AGT1* promoter drove striking, rapid, and unusual ploidy evolution in a wild yeast. Our results thus provide one of the clearest mechanistic insights into the basis of a ploidy fitness effect in fungi.

How generalizable might these principles be? Given the evolutionary lability of ploidy, its link to cell type in many fungal species, and evidence for interactions between cell type and conditionally adaptive traits in other fungal systems, we propose that environment- and genotype-specific regulatory nuances might play a broad role in shaping both the extant diversity of fungal ploidy states and the conflicting, and often cryptic, ploidy and cell type evolution seen in systems experiencing intense selection. This view argues that interactions between cell types, ploidy states, and conditionally adaptive traits may be common during fungal evolution and may influence fungal life cycles more than is currently appreciated.

## DATA AVAILABILITY STATEMENT

Strains and plasmids are available upon request. All raw sequencing data has been deposited at NCBI SRA under BioProject PRJNA894214.

## ACKNOWLEDGMENTS

We thank Kyle Nishikawa and Vatsan Raman for access to and assistance with their flow cytometer and Nathaniel Sharp and members of the Hittinger Lab for constructive feedback. This material is based upon work supported by the National Science Foundation Graduate Research Fellowship Program under Grant No. DGE-1747503, the National Science Foundation under Grant Nos. DEB-1442148 and DEB-2110403, the USDA National Institute of Food and Agriculture (Hatch Project 1020204), and in part by the DOE Great Lakes Bioenergy Research Center (DOE BER Office of Science DE–SC0018409). Any opinions, findings, and conclusions or recommendations expressed in this material are those of the author(s) and do not necessarily reflect the views of the National Science Foundation. Research in the Hittinger Lab is supported an H. I. Romnes Faculty Fellowship, supported by the Office of the Vice Chancellor for Research and Graduate Education with funding from the Wisconsin Alumni Research Foundation. The work conducted by the U.S. Department of Energy Joint Genome Institute, a DOE Office of Science User Facility, is supported under Contract No. DE-AC02-05CH11231. JGC was also supported by a Predoctoral Training Grant in Genetics funded by the National Institutes of Health under Grant No. 5T32GM007133. KJF was a Morgridge Metabolism Interdisciplinary Fellow supported by the Morgridge Institute for Research Metabolism Theme. The funders had no role in study design, data collection and analysis, decision to publish, or preparation of the manuscript.

## Author contributions

JGC: Conceptualization, Funding Acquisition, Formal Analysis, Investigation, Writing—Original Draft, Writing—Review and Editing

KJF: Investigation

TKS: Resources

CTH: Conceptualization, Funding Acquisition, Writing—Review and Editing

---

**Figure S1.**
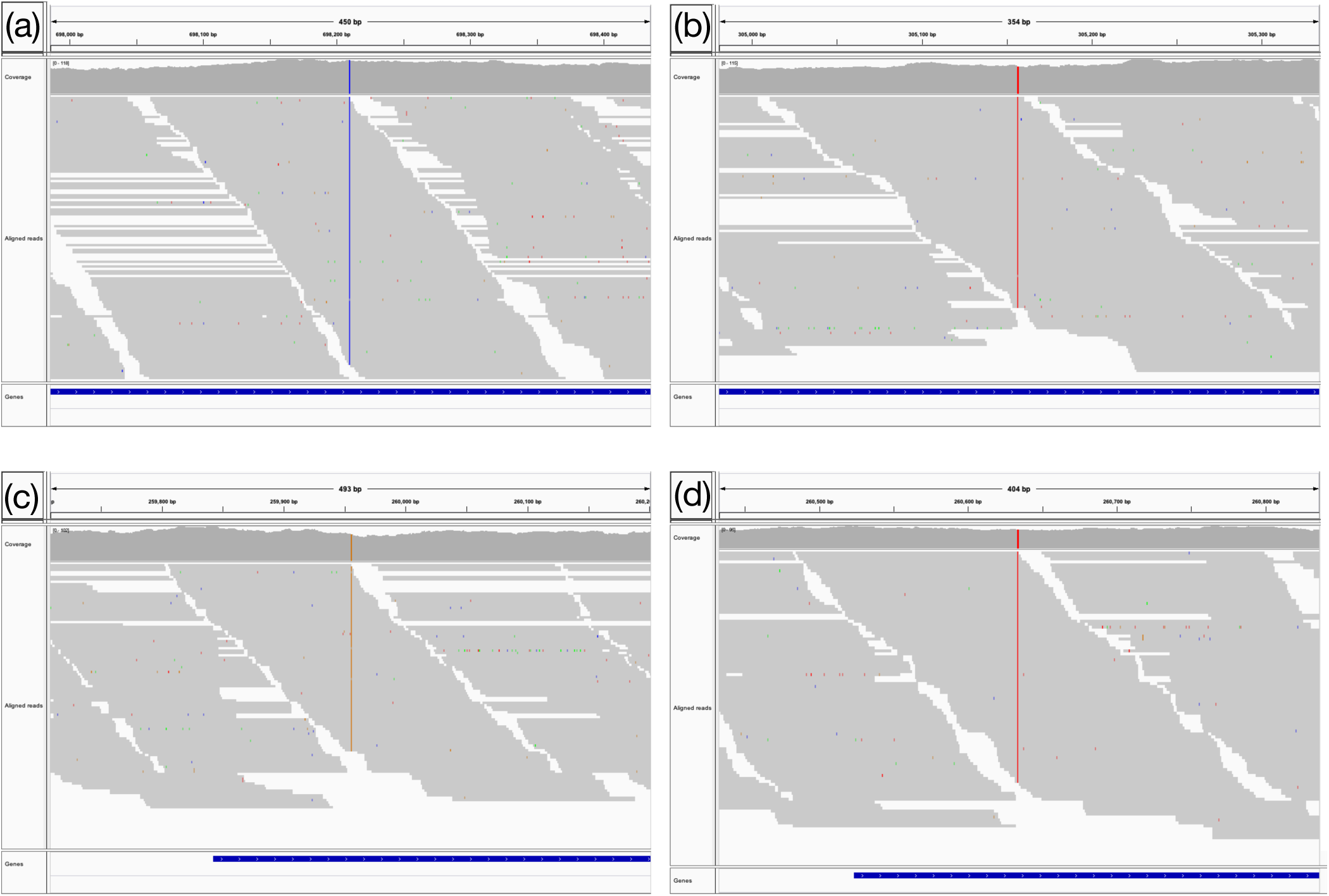
Evolved clones possess only a single allele at each variable site. Genome browser tracks showing aligned Illumina reads from the population 1 clone at *SIR4* (a), *IRA1* (b), and *YDJ1* (c), and from the population 2 clone at *LAM5* (d). Mapped reads are depicted as grey bars with mismatches colored according to base identity.

**Figure S2.**
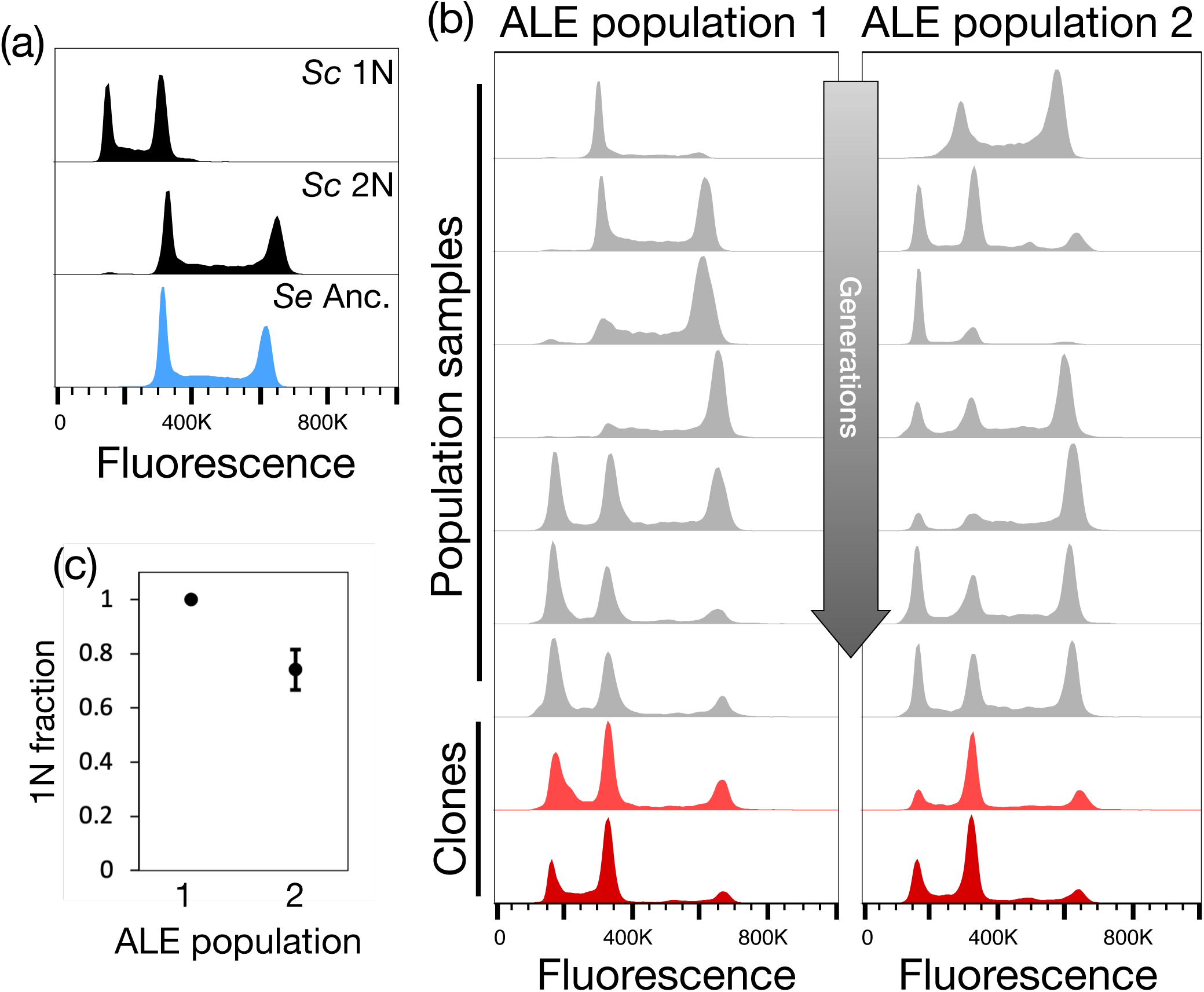
Ploidy variation across the adaptive evolution experiment. (a) Smoothed histograms of cellular DNA content for asynchronous haploid (top panel) and diploid (middle panel) *S. cerevisiae* (*Sc*) controls and the wild-type *S. eubayanus* (*Se* Anc.) strain (bottom panel, reproduced from the same data as in Figure 1). (b) Histograms for population-level samples from both adaptive laboratory evolution (ALE) replicates (grey) and clonal isolates from each population (biological replicates in red). The bottom panel for each clone (dark red) is reproduced from the same data as in Figure 1. For population traces, panels are arranged from top to bottom with increasing time and number of ALE generations. The bottom traces represent the terminal timepoint from which the adapted clones were isolated and from which we quantitatively assessed haploid frequency. (c) Fraction of haploids in the terminal timepoint of each ALE population assayed by *MAT* locus PCR genotyping. Points and bars show the mean and standard error of four experiments.

**Figure S3.**
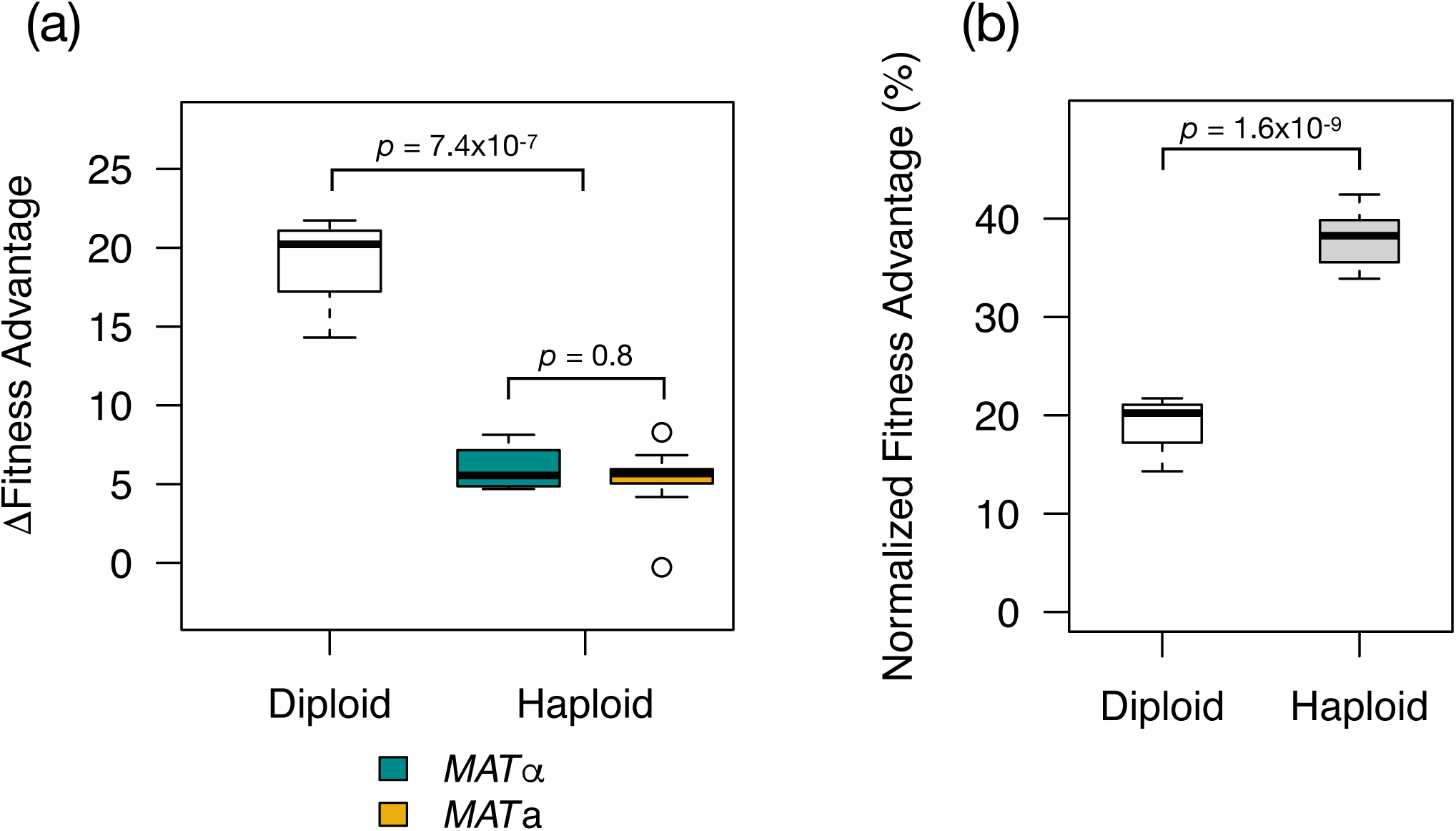
Increased expression of *AGT1* is adaptive. (a) Boxplots show the differences in fitness of diploids and haploids with an extra copy of *AGT1*, compared to the respective parent strain. While haploids experience a smaller change in fitness than diploids, the overall fitness of haploids with increased *AGT1* expression is significantly and substantially higher than that of diploids with increased *AGT1* expression (b).

**Figure S4.**
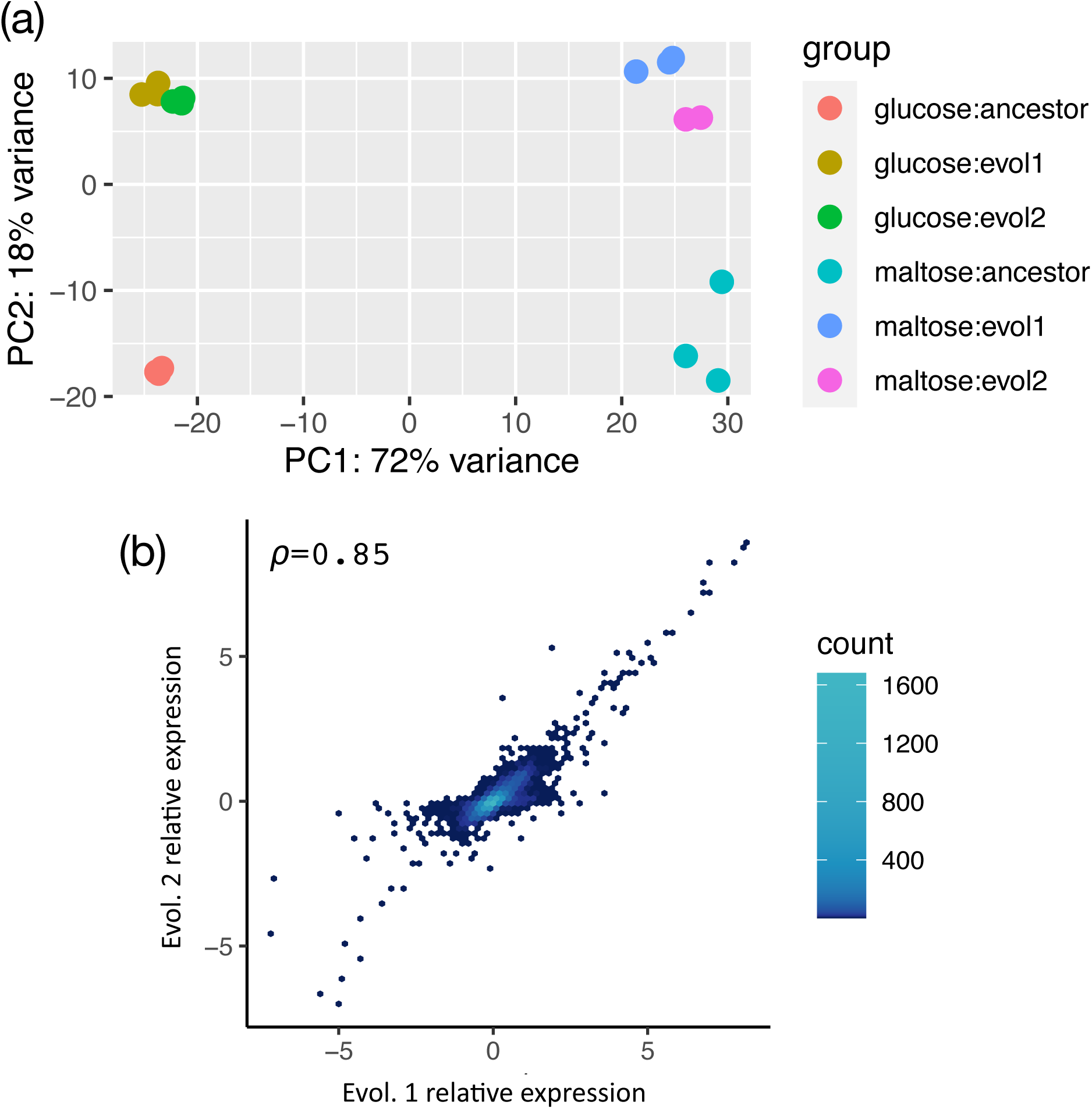
Transcriptomes of the independently evolved haploids are similar. (a) Principal component analysis (PCA) of normalized gene expression for the mRNA-seq libraries used here. Points represent individual libraries, colored by strain and growth condition. Carbon source (PC1) describes a large majority of the variance in the data; PC2 partitions along genotype with evolved haploids (evol1, 2) being distinct from the wild-type diploid (ancestor). (b) Scatterplot of the relative expression of all genes in both conditions for each evolved haploid with hexbin color indicating the density of points. Pearson’s *ρ* is given inset (*p* < 2.2×10^-16^).

**Figure S5.**
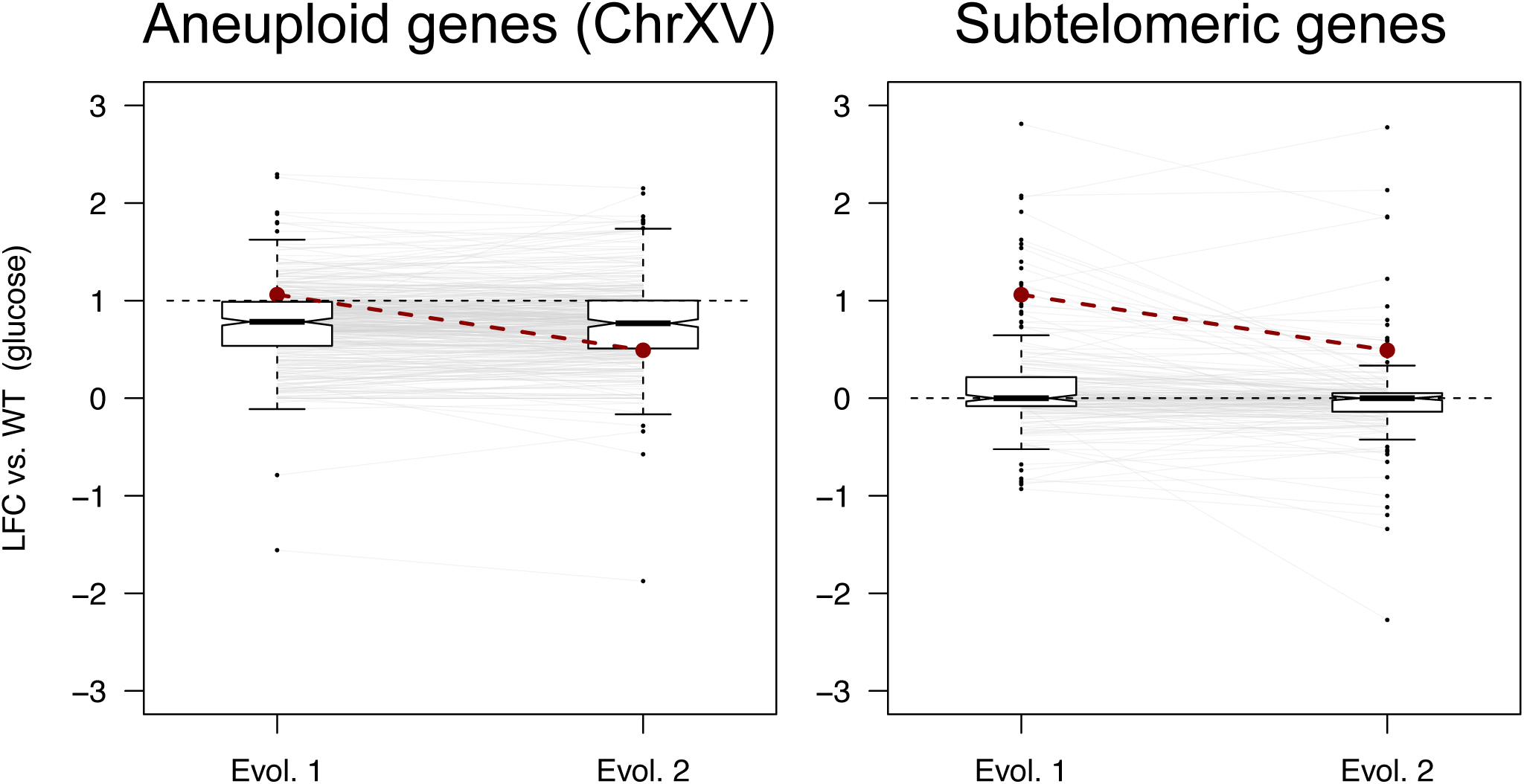
As in Fig. 4, boxplots show log_2_-transformed fold changes (LFC) of gene expression on glucose in evolved haploids compared to the wild-type strain for genes on aneuploid ChrXV (n=370) and subtelomeric genes (n=200). Whiskers extend to 1.5X the interquartile range. Lines connect the y-axis coordinates of the same gene in each evolved isolate; axes are scaled such that an occasional outlier is truncated from the plot space for a single strain. *AGT1* expression is plotted as red dots and lines, and black dashed lines indicate the null expectation for expression values.

**Table S1.** Strains and plasmids used in this study.

**Table S2.** Oligonucleotides used in this study.

**Table S3.** Mutations in evolved isolates.

**Table S4.** Full differential expression analysis results.

